# Development of a dual fluorescent reporter system in *Clostridioides difficile* reveals a division of labor between virulence and transmission gene expression

**DOI:** 10.1101/2022.03.03.482933

**Authors:** M. Lauren Donnelly, Shailab Shrestha, John Ribis, Pola Kuhn, Maria Krasilnikov, Carolina Alves Feliciano, Aimee Shen

## Abstract

The bacterial pathogen *Clostridioides difficile* causes gastroenteritis through its production of toxins and transmits disease through its production of resistant spores. Toxin and spore production are energy-expensive processes that are regulated by multiple transcription factors in response to many nutritional inputs. While toxin and sporulation genes are both heterogeneously expressed in only a subset of *C. difficile* cells, the relationship between these two sub-populations remains unclear. To address whether *C. difficile* coordinates the generation of these sub-populations, we developed a dual transcriptional reporter system that allows toxin and sporulation gene expression to be simultaneously visualized at the single-cell level using chromosomally-encoded mScarlet and mNeonGreen fluorescent transcriptional reporters. We then adapted an automated image analysis pipeline to quantify toxin and sporulation gene expression in thousands of individual cells in different media conditions and genetic backgrounds. These analyses revealed that toxin and sporulation gene expression rarely overlap during growth on agar plates, but broth culture increases this overlap in a manner dependent on the multifunctional RstA transcriptional regulator. Our results suggest that certain growth conditions promote a “division of labor” between transmission and virulence gene expression, highlighting how these subpopulations are influenced by environmental inputs. Given that recent work has revealed population-wide heterogeneity for numerous cellular processes in *C. difficile*, we anticipate that our dual reporter system will be broadly useful for determining the overlap in these subpopulations.

**IMPORTANCE:** *Clostridioides difficile* is an important nosocomial pathogen that causes severe diarrhea by producing toxins and is transmitted by producing spores. While both processes are crucial for *C. difficile* disease, only a subset of cells express toxins and/or undergo sporulation. Whether *C. difficile* coordinates the relationship between these energy-expensive processes remains unknown. We developed a dual fluorescent reporter system coupled with an automated image analysis pipeline to rapidly characterize expression two genes of interest across thousands of bacterial cells. Using this reporter system, we discovered that toxin and sporulation gene expression appear to undergo a “division of labor” in certain growth conditions, particularly during growth on agar plates. Since *C. difficile* specializes into subpopulations for numerous vital cellular processes, this novel dual reporter system will enable future studies aimed at understanding how *C. difficile* coordinates various subpopulations throughout its infectious disease cycle.

## INTRODUCTION

*Clostridioides difficile* is a Gram-positive, spore-forming, anaerobic pathogen that is the leading cause of nosocomial diarrhea worldwide and the most common healthcare-associated infection in the United States (1, 2). *C. difficile* infections typically occur in patients whose normal colonic microflora is disrupted such as individuals who have undergone antimicrobial therapy (1). *C. difficile* infections often disseminate in healthcare settings because its resistant, infectious, aerotolerant spore morphotype is transmitted by the fecal-oral route and can persist in the environment for long periods (3–6). When *C. difficile* spores are ingested, they germinate in the gut of dysbiotic hosts in response to specific bile acids (7) and outgrow into vegetative cells. These cells produce the glucosylating toxins, Toxin A (TcdA) and Toxin B (TcdB), that are responsible for *C. difficile* disease symptoms (8). By glucosylating Rho family GTPases, the toxins induce actin cytoskeleton collapse and loss of tight junctions in the colon (8). The resulting damage to the colonic epithelium elicits a massive host inflammatory response, which can lead to pseudomembranous colitis, colonic perforation, toxic megacolon, and even death (9).

Toxin production and spore formation are frequently coordinated in spore-forming bacteria, with toxin production being induced during early stages of sporulation in pathogens like *Clostridium perfringens* and some strains of *C. botulinum* and *Bacillus thuringiensis* (10–15). This temporal order likely enhances the transmission of these pathogens because the toxins induce diarrhea in or death of the host to promote dissemination of the spores into the environment (10, 16). However, the relationship between toxin and sporulation genes is often strain-dependent. For example, some *B. thuringiensis* strains generate a “division of labor” in which some cells within the population express toxin genes and a different subset induces sporulation (16, 17). The relationship between toxin and sporulation gene expression in *C. difficile* has not been thoroughly analyzed, although recent work has shown that both toxin and sporulation genes are heterogeneously expressed within a given population of *C. difficile* cells, with only a subset of cells expressing toxin genes and a subset expressing sporulation genes. While these two subpopulations can overlap (18), it is unclear how frequently this occurs on a population-wide level.

Numerous environmental signals control toxin and sporulation gene expression in *C. difficile*. For example, both sets of genes are repressed by glucose and preferentially induced during stationary phase in *C. difficile*, suggesting that nutrient starvation induces these processes (19). The levels of c-di-GMP, autoinducing peptides, and other signaling molecules also modulate these processes (20, 21). Notably, a complex network of overlapping genetic circuits control toxin and sporulation gene expression in *C. difficile*. *tcdA* and *tcdB* toxin gene expression is activated by TcdR, an alternative sigma factor whose levels determine toxin gene bistability (18). TcdR auto-regulates its production through a positive feedback loop, but several transcription factors regulate *tcdR* expression in response to nutrient availability and growth phase (22). Sporulation is controlled by the master transcriptional activator, Spo0A, which initiates sporulation by inducing the expression of genes encoding sporulation-specific sigma factors that subsequently drive sporulation (23). *spo0A* transcription and/or Spo0A activity are controlled by several of the same regulatory factors that affect toxin production (20, 24–26). For example, CodY is a global regulator that represses both sporulation and toxin gene expression (27) when branched chain amino acids and GTP are abundant (27–30). In addition, the carbon catabolite protein A (CcpA) represses toxin and sporulation gene expression in the presence of glucose and other carbohydrates (31). SigH is a positive regulator of sporulation and a negative regulator of toxin production (32), while the bifunctional transcription factor, RstA, represses toxin production but enhances sporulation. RstA directly inhibits transcription of *tcdA, tcdB*, and *tcdR* (33), while indirectly promoting the expression sporulation genes (34).

Interestingly, some studies suggest that toxin and sporulation gene expression may be coordinated in *C. difficile* because cross-talk between TcdR and Spo0A function has been reported (35–37). For example, Spo0A negatively regulates toxin production in some but not all isolates of *C. difficile* likely through indirect mechanisms (36), and loss of toxin production due to mutation of *tcdR* enhances sporulation in some strain backgrounds (35). Notably, many of these findings are strain-specific as well as highly dependent on growth and media conditions, leading to conflicting reports of the relationship between sporulation and toxin production (38).

Disentangling how *C. difficile* coordinates toxin and sporulation gene expression requires the development of transcriptional reporters that can be simultaneously visualized in individual cells. In this study, we optimized a chromosomally-encoded dual reporter system for use in *C. difficile* that overcomes *C. difficile*’s high intrinsic autofluorescence (39). Using these reporters, we addressed whether *C. difficile* coordinates toxin and sporulation gene expression in different media conditions and mutant backgrounds. Our results suggest that certain growth conditions promote a “division of labor” between toxin and sporulation gene expression, with minimal overlap between the subsets of cells expressing toxin and sporulation genes. Given that recent studies have identified several genes beyond toxin and sporulation genes with bimodal patterns of expression (40–42), our novel dual reporter system should be broadly useful for studying how these different sub-populations of *C. difficile* cells overlap and how nutritional and environmental inputs affect this overlap.

## RESULTS

### Development of chromosomally-encoded dual-color transcriptional reporters for use in *C. difficile*

To visualize toxin and sporulation gene expression simultaneously, we first needed to identify reporters with non-overlapping spectra that were also bright enough to overcome *C. difficile*’s autofluorescence in the green channel (39, 43). This autofluorescence has traditionally limited the utility of green-emitting reporters like GFP and phiLOV, an FMN-based reporter to genes that are highly expressed (39, 44). However, mNeonGreen (**mNG**) is a rapidly maturing, yellow-green monomeric fluorescent protein made by *Branchiostoma lanceolatum* that is ∼4-fold brighter than GFP (45, 46), so we hypothesized that it might be bright enough to use as a fluorescent transcriptional reporter. Another advantage of mNeonGreen is that its spectral properties are closer to YFP, which reduces some of the autofluorescent signal from *C. difficile*. Finally, mNeonGreen is spectrally compatible with mScarlet (**mSc**), a derivative of mCherry that is ∼3-fold brighter than mCherry (47), such that mNeonGreen and mScarlet can be used in dual transcriptional reporter and protein localization systems(46, 48).

To test whether mNeonGreen and/or mScarlet fluorescent reporters would be bright enough for use in *C. difficile*, we codon-optimized *mNeonGreen* and *mScarlet* and coupled these genes to two different constitutive promoters: P*slpA* and P*cwp2*. *slpA* encodes the major S layer protein, and is the most highly transcribed gene in our prior RNA-Seq analyses in *C. difficile* (49–51), while *cwp2* encodes a cell wall protein (52) and its promoter has previously been used as a constitutive promoter(53).The promoters were followed by the ribosome binding site of the *slpA* gene, and the resulting constructs were integrated in single-copy into the *pyrE* locus of 630Δ*erm*Δ*pyrE* using allele-coupled exchange (44). After the resulting strains were grown overnight in TY media, we spotted the cultures onto agarose pads under ambient conditions so that the oxygen would allow the mNeonGreen and mScarlet fluorophores to mature. Fluorescent signal above background was observed in all cells for both the *mNeonGreen* (Fig 1A) and *mScarlet* constitutive reporters (Fig 2A).

**Figure 1.**
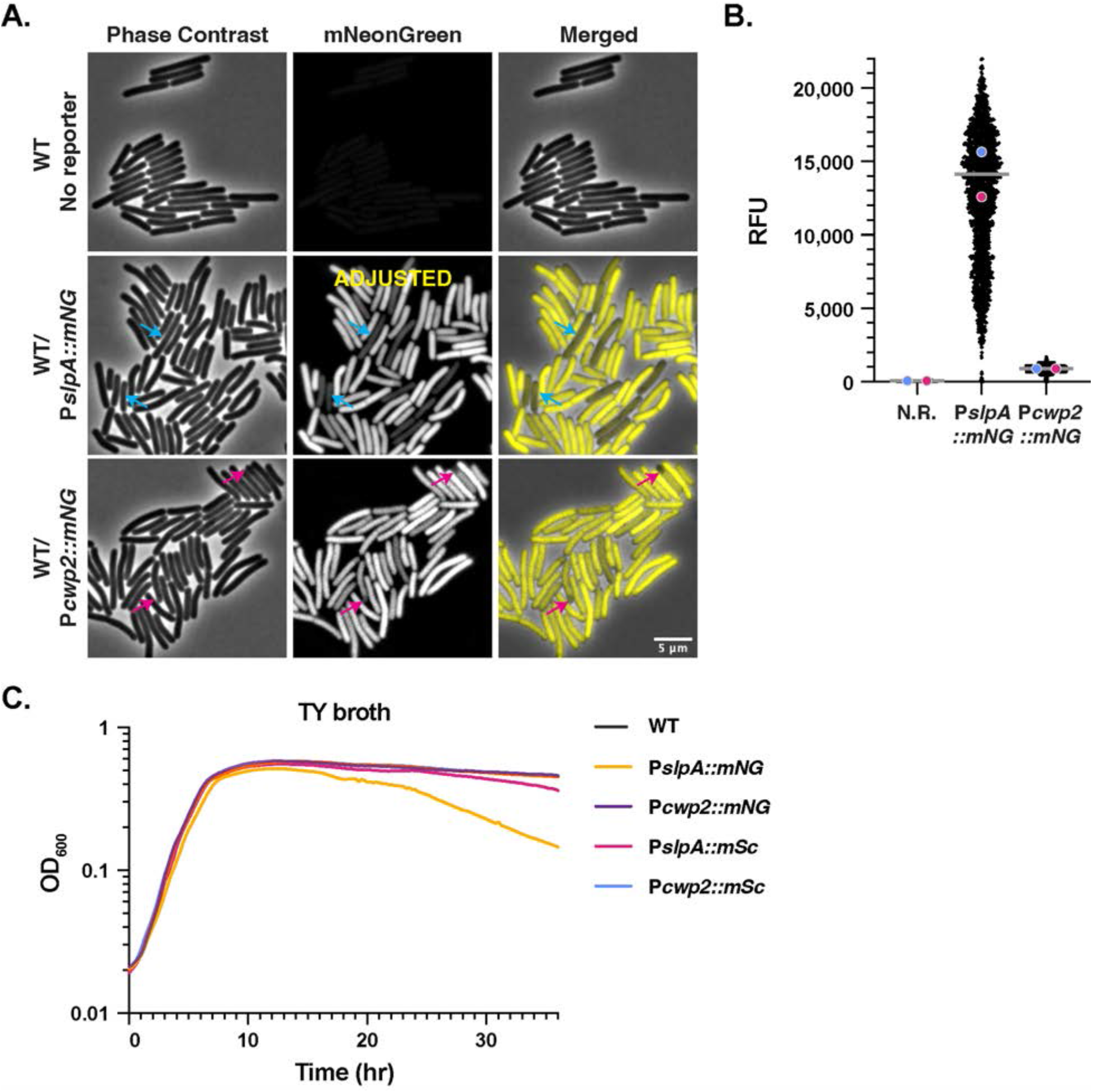
Identification of constitutive *mNeonGreen* reporters that overcome *C. difficile’s* autofluorescence. (A) Fluorescence microscopy analyses of live bacterial cells showing wild type 630Δ*erm* (no reporter) and 630Δ*erm* carrying *mNeonGreen* coupled to either the *slpA* or *cwp2* promoters after overnight growth in TY media. Phase-contrast microscopy was used to visualize the cells. Blue arrows highlight cells where the mNeonGreen signal appears reduced relative to other cells in the population for the P*slpA::mNG* strain. Pink arrows highlight lower levels of mNeonGreen signal in the forespore of cells undergoing sporulation. The merge of phase-contrast and mNeonGreen pseudo-colored in yellow images is shown. (B) SuperSegger-based quantification (55) of the mean fluorescent intensity for each cell is shown as a black dot on the scatterplot. The magenta and blue dots represent the median fluorescent intensity for the first and second biological replicates, respectively. The gray line represents the mean fluorescence value for each reporter based on the average of the two biological replicate’s median value (79). N.R. indicates wild type with no reporter. (C) OD_600_ growth curve of the indicated strains during growth in TY broth. The graph shown is a single biological replicate that is representative of three biological replicates.

**Figure 2.**
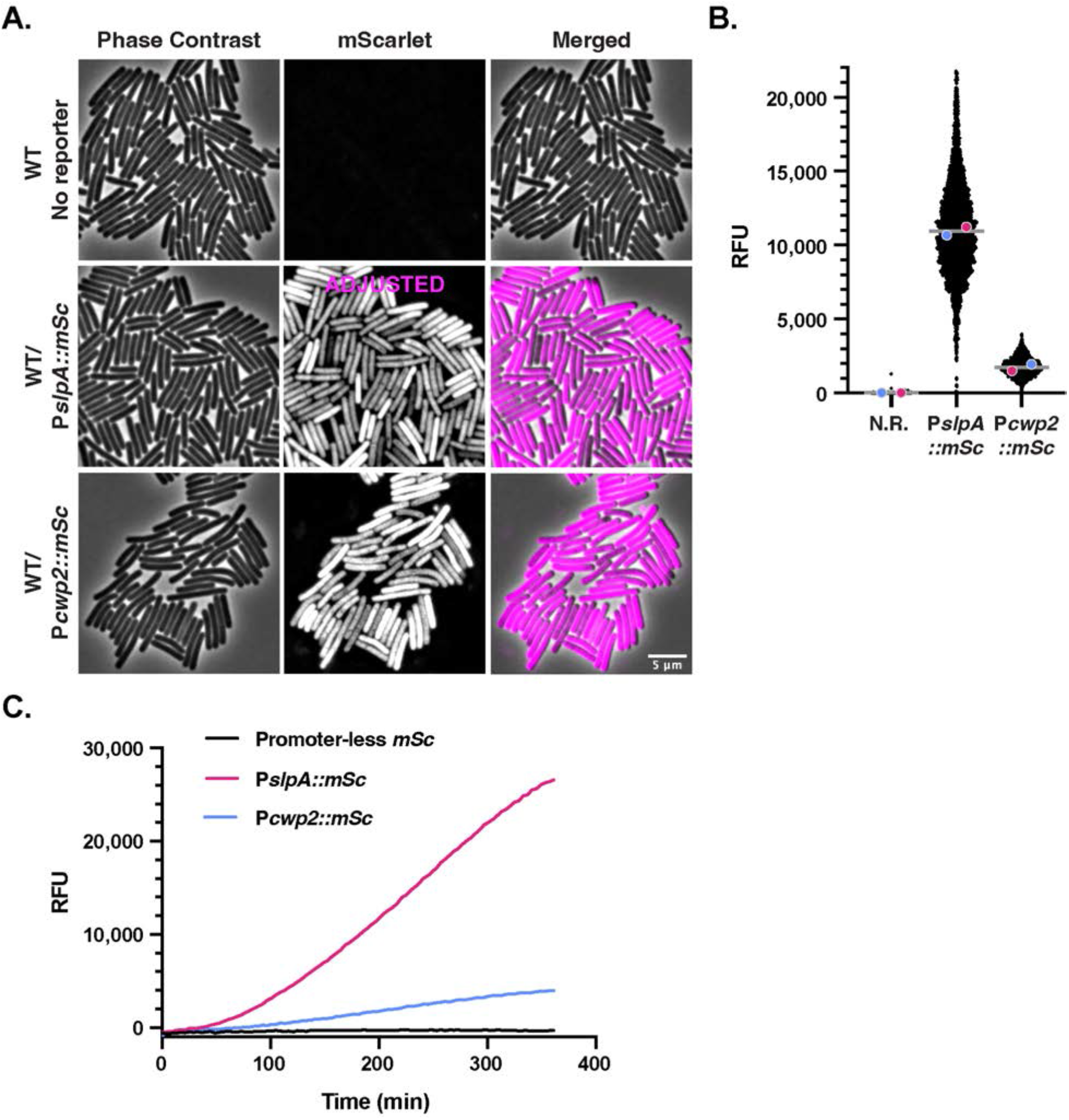
Optimization of constitutive mScarlet reporters for use in *C. difficile*. (A) Fluorescence microscopy analyses of fixed cells for wild type 630Δ*erm* (no reporter) and 630Δ*erm* carrying *mScarlet* coupled to either the *slpA* or *cwp2* constitutive promoters after overnight growth in TY broth. Phase-contrast microscopy was used to visualize all bacterial cells. The merge of phase-contrast and mScarlet pseudo-colored in magenta is shown. The P*slpA::mSc* signal was adjusted for brightness/contrast because this reporter is so much brighter than P*cwp2::mSc*. (B) SuperSegger-based quantification of the mean fluorescent intensity for each cell is shown as a black dot on the scatterplot. The magenta and blue dots represent the median fluorescent intensity for the first and second biological replicates, respectively. The gray line represents the mean fluorescence value for each reporter based on the average of the two biological replicate’s median value (79). N.R. indicates wild type with no reporter. (C) Fluorescent intensities of overnight TY cultures of the indicated strains after exposure to oxygen over the course of 36 hours.

To quantify the fluorescent signal at the single-cell level, we adapted SuperSegger, a machine learning-based cell image analysis suite designed for time-lapse microscopy data (55), to calculate the mean fluorescent signal intensity for each cell in still images. SuperSegger readily identifies individual cells even if they are in close proximity, enabling automated quantification of thousands of individual cells. Using SuperSegger, we determined that the fluorescent signal was brighter for both the P*slpA::mNG* and P*slpA::mSc* reporters relative to the analogous P*cwp2* reporters (Fig 1B & 2B). Since the chromosomally-encoded mNeonGreen reporters were bright enough to visualize above *C. difficile*’s autofluorescence, *mNeonGreen* and *mScarlet* could be suitable for use in chromosomally-encoded dual transcriptional reporter systems. Importantly, chromosomal expression helps avoid issues related to variable plasmid-copy number and plasmid segregation when examining the activity of gene-specific promoters at the single-cell level (39).

### mScarlet and mNeonGreen have distinct benefits and drawbacks for use in *C. difficile*

To better understand the utility of the *mScarlet* and *mNeonGreen* reporters for such applications, we explored the advantages and disadvantages associated with of the two reporters. This analysis was prompted by our finding that two populations of P*slpA::mNG* cells could be seen when their fluorescence intensities were plotted as a histogram (< 12,000 RFUs, **Fig S1**), whereas P*cwp2::mNG* and the *mSc* constitutive reporter strains exhibited a largely normal distribution (**Fig S1**). We hypothesized that P*slpA::mNG* cells with lower signal intensities (Fig 1B) may be experiencing toxicity related to high levels of mNeonGreen, since cells with lower mNeonGreen levels also stain less intensely with the Hoechst nucleoid stain (Fig 1, blue arrows), which may indicate decreased viability of this subset of cells. Since a recent report showed that *C. difficile* expressing high levels of the *mCherry* reporter have reduced viability (39), we compared the growth of the *mNG* and *mSc* reporter strains in TY media relative to WT (no reporter). The optical density of the P*slpA::mNG* strain, but not the other strains (Fig 1C), decreased after ∼15 hrs of growth. Given that the P*slpA::mNG* strain produces considerably more mNeonGreen based on fluorescence measurements relative to P*cwp2::mNG* (Fig 1B), this data suggest that high levels of mNeonGreen protein are deleterious to the cell during late stationary phase.

Since recent work has shown that *C. difficile* is more susceptible to autolysis during growth in BHIS vs. TY media (56), we analyzed the behavior of our constitutive *mNG* and *mSc* reporter strains after culturing them overnight in BHIS broth. In contrast with the relatively uniform fluorescence of the reporter strains during overnight growth in TY broth (Fig 1A), only a small fraction of P*slpA::mNG* and P*cwp2::mNG* cells were fluorescent following overnight growth in BHIS broth (**Fig S2A,B**). This result suggests that either mNeonGreen is less stable in late stationary phase cells following growth in this medium or cells producing mNeonGreen lose their viability at this timepoint. Consistent with the latter hypothesis, the P*slpA*::*mNG* strain exhibited an even more severe decrease in optical density relative to the other strains during late stationary phase growth in BHIS broth, and its growth in exponential phase lagged behind the other strains in BHIS media (**Fig S2C**). This issue was specific to *mNG* because the P*slpA::mSc* and P*cwp2::mSc* strains retained their fluorescence even in BHIS overnight cultures (**Fig S2B**). A similar loss in mNeonGreen signal was observed with the P*slpA::mNG* reporter when grown in BHIS regardless of whether the cells were fixed vs. live cells or grown on solid agar or in broth culture (data not shown). Taken together, these data suggest high levels of mNeonGreen may lead to autolysis in *C. difficile* (56).

While high level mScarlet production did not appear to strongly affect the growth of *C. difficile*, we noticed that the fluorescence of mScarlet-producing strains increased over time following exposure to oxygen relative to mNeonGreen-producing strains. This finding is consistent with mScarlet’s long maturation time of 174 min at 37°C (47) relative to mNeonGreen’s maturation time of 10 min at 37°C, (45). To analyze the kinetics of mScarlet’s maturation in *C. difficile* cells upon exposure to oxygen, we measured its bulk fluorescent signal over time after exposing broth cultures of mScarlet-producing strains to ambient oxygen using a plate reader (Fig 2C). The mScarlet signal started to peak around 6 hrs after fixation and exposure to oxygen, highlighting the importance of extended oxygen exposure for visualizing mScarlet. While mScarlet fluorescence was detectable in live cells even 1 hour after exposure to oxygen, we found that waiting 2 hours markedly increased the fluorescence level. Since oxygen exposures >2 hrs resulted in chromosome fragmentation, we fixed samples containing the mScarlet fluorescent reporter using previously established procedures (43). Notably, no difference in mNeonGreen fluorescence was observed between live and fixed cells, consistent with mNeonGreen’s fast maturation time (45).

Despite mScarlet requiring fixation for maximal detection in *C. difficile*, we found that the mScarlet is more sensitive than mNeonGreen because *C. difficile* has minimal autofluorescence in the red channel, and its production appears less deleterious to *C. difficile* cells in late stationary phase. Conversely, mNeonGreen is better suited for imaging live cells (i.e. without fixation) due to its short maturation time, although it can cause toxicity to *C. difficile* in certain growth conditions when produced in large amounts (Figs 1C and **S1C**).

### Constitutive reporter fluorescence is reduced in sporulating cells especially in the forespore, but forespore-specific reporters can be generated

While analyzing constitutive reporter strains in different growth media, we observed that the fluorescence of the reporters appeared to decrease in the forespore region of cells undergoing sporulation (Fig 1, pink arrows and Fig 3, yellow arrows). The decreased expression of genes encoding surface proteins like SlpA and Cwp2 in the forespore might be expected because the mature spore does not produce these surface proteins (59). However, it is possible that the reduced signal for the constitutive reporters is an artifact caused by the inherent instability of the fluorescent proteins in the forespore. To address this possibility, we constructed *mNG* and *mSc* reporters driven by the forespore-specific *sspB* promoter (P*sspB::mNG* and P*sspB::mSc*) (49, 60). *sspB* encodes a small acid-soluble spore protein (SASP) that helps protect spore DNA from UV damage (61) and whose expression is σ^F^- and σ^G^-dependent and thus localized to the forespore (49, 60). Fluorescence in the P*sspB::mSc* reporter strain was localized to the forespore, indicating that it is possible for mScarlet to be stably produced in the forespore (Fig 3). Surprisingly, the mScarlet signal was also occasionally observed throughout the cytosol of rod-shaped cells with no visible signs of sporulation based on their morphology (i.e. following asymmetric division, Fig 3, orange arrows, (57)). Since this apparent vegetative cell signal was not observed when we integrated the reporter into a Δ*spo0A* mutant (Fig 3), which cannot initiate sporulation (37), the fluorescent signal likely originates in cells that have induced sporulation but have not completed asymmetric division and are thus predivisional cells (62). While the aberrant activation of σ^F^ in predivisional cells has been reported in *B. subtilis* at low frequency (0.5%, (62)), we note that it is also possible that the mScarlet signal may have leaked out of the forespore, since the predivisional cells exhibit high mScarlet fluorescence but lack Hoechst staining of their nucleoid region (Fig 3, green arrows). Regardless of the origins of this signal, we note that a similar “predivisional” signal is observed for additional forespore-specific *mScarlet* reporters that we tested involving fusions to the *spoIIQ*, *pdaA*, and *sspA* promoters (data not shown).

**Figure 3.**
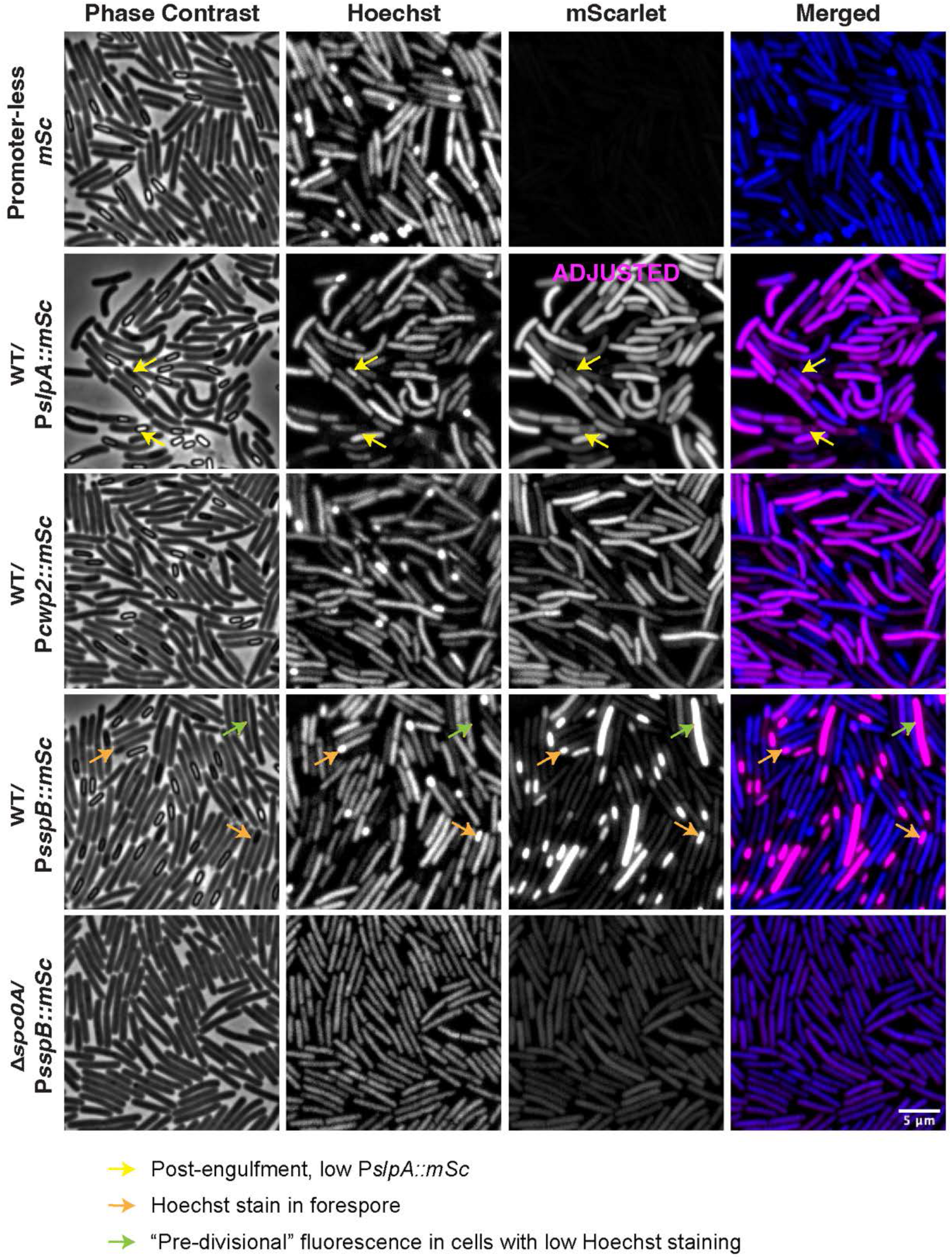
Expression levels of constitutive genes decrease in sporulating cells particularly in the forespore, but forespore-specific gene expression can still be visualized. (A) Fluorescent microscopy analyses of P*slpA::mSc,* P*cwp2::mSc*, and P*sspB::mSc* reporter strains. Phase-contrast microscopy was used to visualize sporulating cells (Phase). Hoechst staining was used to visualize the nucleoid. The merge of the Hoechst (blue) and the mScarlet signal (magenta) is shown. Reduced mScarlet signal in the forespore of cells that are visibly sporulating is highlighted with a yellow arrow in the P*slpA::mSc* strain, but this was also observed for the P*cwp2::mSc* strain. Cells that are visibly sporulating based on the concentration of the Hoescht stain in the forespore is shown in orange arrows. Bright P*sspB::mScarlet* signal in cells that are either pre-divisional or sporulating cells where the forespore has become compromised and mScarlet has become distributed across the entire cell length in a Spo0A-dependent manner are highlighted by green arrows. Images are representative of three biological replicates. Microscopy was performed on live sporulating cultures 15-18 hr after plating on SMC sporulation agar. P*slpA::mSc* is displayed with adjusted settings to accommodate its bright signal relative to P*cwp2* and P*sspB*.

While we readily detected the mScarlet signal in the forespore of sporulating P*sspB::mSc* cells, when we coupled the *sspB* promoter to *mNG*, mNeonGreen fluorescence in the forespore was barely above background autofluorescence (**Fig S3**, orange arrows), highlighting how *C. difficile*’s intrinsic autofluorescence reduces the sensitivity of *mNeonGreen*-based transcriptional reporters. Regardless, these experiments demonstrate that mNeonGreen and mScarlet can still be used to detect transcription in the forespore, implying that the reduced signal for P*slpA-* and P*cwp2-*driven reporters in the forespore and mother cell of sporulating cells likely reflects a general downregulation in gene expression as the forespore matures and the mother cell prepares to lyse and release the spore.

### Toxin gene expression appears elevated in strains with decreased sporulation

Having established the utility of *mNeonGreen* and *mScarlet* transcriptional reporters for visualizing gene expression when chromosomally expressed in *C. difficile*, we next sought to investigate the relationship between toxin and sporulation gene expression at the single-cell level. We first generated chromosomal *mNG* and *mSc* reporters coupled to the *tcdA* promoter (P*tcdA::mNG* and P*tcdA::mSc*, respectively) to visualize the sub-population of cells that express toxin genes. These constructs use the same promoter sequence as the plasmid-based P*tcdA::mCherry* transcriptional reporter previously described (18). As a control, we integrated the P*tcdA::mNG* and P*tcdA::mSc* into the *pyrE* locus of a clean *tcdR* deletion strain because the TcdR alternative sigma factor activates *tcdA* expression (18). We also sought to assess how sporulation impacts toxin expression at the single-cell level by integrating the P*tcdA::mNG* and P*tcdA::mSc* reporters into the *pyrE* locus of a previously constructed Δ*spo0A* mutant because prior work had suggested that Spo0A negatively regulates toxin production in some *C. difficile* strains (36) while others suggest the opposite (63). To further explore the relationship between sporulation and toxin expression, we also moved the P*tcdA* reporter into a strain lacking the bifunctional transcriptional regulator, RstA, which represses toxin production and promotes sporulation (64). While previous work had shown that *ΔrstA* over-expresses toxin genes in bulk population measurements, it was unclear whether loss of RstA impacts the frequency of cells expressing toxin genes and/or the magnitude of their toxin gene expression (64).

We validated our *ΔtcdR* and Δ*rstA* strains by complementing the respective mutants (no reporter) with a wild-type copy of *tcdR* or *rstA* driven by their native promoter expressed from the *pyrE* locus. Importantly, the Δ*tcdR*/*tcdR* complementation strain produced wild-type levels of TcdA in Western blot analyses of overnight TY broth cultures, while no TcdA was detected in the parental Δ*tcdR* strain (**Fig S4A**). As expected, the Δ*rstA* mutant over-produced TcdA (∼4.5-fold more, p < 0.0001), while the complementation strain produced wild-type levels of TcdA. Toxin levels were also elevated (∼4-fold, p < 0.001) in the Δ*spo0A* mutant, while the complementation produced wild-type levels of TcdA.

When P*tcdA::mNG* and P*tcdA::mSc* were integrated into the *pyrE* locus of wild-type 630Δ*erm*, the reporters exhibited a heterogeneous expression pattern within the population of cells (Fig 4A **& S5**) similar to previously described analyses using a plasmid-based reporter (18). Some cells were clearly in a “Toxin ON” state, with bright mNeonGreen or mScarlet fluorescence visible, and other cells lacked detectable fluorescence and were thus “Toxin OFF” (18). As expected, little signal was observed in the Δ*tcdR* mutant background. We also attempted to visualize *tcdB* toxin gene expression using *mNeonGreen* and *mScarlet* reporters in *C. difficile* but both chromosomal and plasmid-based reporters failed to generate fluorescent signal above background.

**Figure 4.**
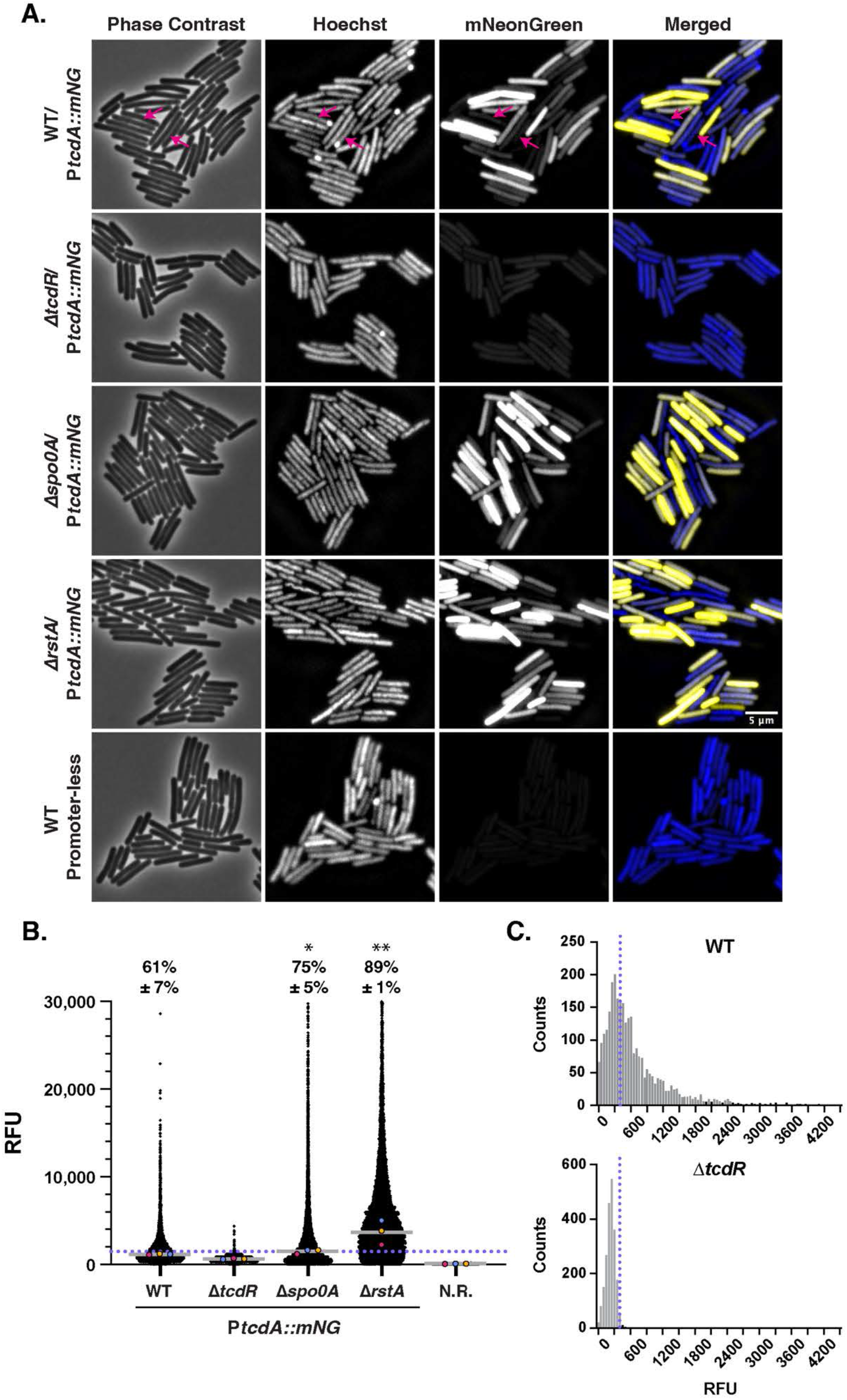
Toxin gene expression is elevated in strains with reduced sporulation (for both mNG and mSc reporters). (A) Fluorescence microscopy analyses of live cells from strains carrying P*tcdA::mNG* toxin gene reporters grown overnight in TY broth relative to a promoter-less *mNG* construct integrated into the *pyrE* locus. Phase-contrast microscopy was used to visualize all bacterial cells, and the nucleoid was stained with Hoechst. The merge of Hoechst (blue) and mNeonGreen pseudo-colored in yellow is shown. Sporulating cells based on Hoescht stain with decreased toxin reporter levels are highlighted with magenta arrows. (B) SuperSegger-based quantification of the mean fluorescent intensity of each cell is shown as a black dot on the scatterplot. Individual cell intensities were quantified from three biological replicates with at least two fields of view per strain per replicate. The magenta, yellow, and blue dots represent the median intensity for the first, second, and third biological replicates, respectively. The gray line represents the mean value of each replicate’s median value. N.R. indicates a strain harboring *mNeonGreen* with no upstream promoter region integrated into the *pyrE* locus. Percentage “Toxin-ON” is displayed ± the standard deviation. “Toxin-ON” cells were calculated using the 99^th^ percentile of the Δ*tcdR* signal as a cutoff (value displayed as a blue dotted line). A minority of points (<1%) are outside the limits of the scatterplot; axes were adjusted to provide an optimal view of the scatterplot trends. Statistical significance was determined relative to wild type using a one-way ANOVA and Tukey’s test. *p < 0.1; ** p < 0.01. (C) Histogram analysis of single-cell fluorescent intensities for the P*tcdA::mNG* reporter in wildtype vs. *ΔtcdR*. The 99^th^ percentile cutoff in Δ*tcdR* used to define cells as “Toxin-ON” is shown by the blue dotted line. Plotted data is a compilation of three biological replicates.

To quantify how many cells were in the “Toxin-ON” state with our P*tcdA* reporters, we set the cut-off for wild-type P*tcdA* expression as above the 99^th^ percentile of the P*tcdA* reporters in the Δ*tcdR* strain background. With this cutoff, 61% of cells in the WT population were “Toxin ON” in the *mNeonGreen* reporter strain (Fig 4B) and 37% were “Toxin ON” in the *mScarlet* reporter strain cells (**Fig S5-S6**). While the fraction of “Toxin ON” cells would appear to be different between the P*tcdA::mNG* and P*tcdA::mSc* reporter strains, we note that slight changes in the cutoff value for “Toxin-ON” cells result in a 5-10% change in the proportion of “Toxin-ON” cells. While the biological significance of a cell that is just under or over the cutoff value set by Δ*tcdR* remains unclear, our single-cell analyses confirm that there is a broad range in toxin reporter expression within wild-type cells.

Our findings with these chromosomal toxin reporters are relatively similar to prior analyses using a plasmid-based P*tcdA::mCherry* reporter, which found that ∼80% of cells are “Toxin ON” in the 630 strain background (18). The slight differences in “Toxin-ON” frequencies between our studies could be due to plasmid copy number variation and/or the greater sensitivity of the plasmid-based reporter due to its multi-copy nature. Regardless, our chromosomal P*tcdA* fluorescent reporters gave a similar distribution of “Toxin-ON” cells as a plasmid-based P*tcdA:: mCherry* reporter (18) when plotted as a histogram (**Fig S7**). Notably, the P*tcdA::mNG* reporter exhibited a similar long-tailed distribution as the P*tcdA::mCherry* plasmid-based reporter in the 630 strain background, in contrast with the bimodal distribution in toxin gene expression observed for some *C. difficile* strains (18).

Notably, the long-tailed distribution in toxin reporter expression was even greater for Δ*spo0A* and Δ*rstA* cells (**Fig S7**). *Δspo0A* exhibited a ∼20% increase in the magnitude of toxin gene expression at the single-cell level, and a ∼10-15% increase in the frequency of “Toxin-ON” cells (75% for *mNG* and 48% for *mSc* reporters relative to WT (Figs 4 **& S5**)). Δ*rstA* cells expressed the toxin reporter at higher levels (∼2-3-fold increase relative to WT), and almost every Δ*rstA* cell was “Toxin-ON” (89% for *mNG* and 78% for *mSc* cells). These results are largely consistent with the elevated TcdA levels observed in Δ*spo0A* and Δ*rstA* mutants (**Fig S4**) and the previous observation that loss of RstA derepresses toxin gene expression in bulk (34, 64). Since loss of either Spo0A or RstA increases the proportion of cells expressing the toxin reporters as well as the magnitude of their expression, and since Δ*spo0A* cannot sporulate and Δ*rstA* sporulates at reduced levels, our results suggest that toxin and sporulation genes might be inversely related.

### Sporulation is not elevated in a toxin-less strain

Consistent with this hypothesis, sporulating cells in these samples (visible based on their Hoechst staining) were frequently observed to be “Toxin-OFF” (Fig 4A**, pink arrows**). To assess whether toxin gene expression decreases sporulation, we visualized *C. difficile* sporulation gene expression in strains that either cannot produce toxin (Δ*tcdR*) or produce elevated levels of toxin (Δ*rstA*). To this end, we coupled the sporulation-specific *sipL* promoter to *mNeonGreen* and *mScarlet* (P*sipL::mNG* and P*sipL::mSc*, respectively). *sipL* is expressed under the control of the mother cell-specific sigma factor σ^E^ (49, 60), so its expression should localize only to the mother cell cytosol and not the forespore. We integrated the P*sipL::mNG* and P*sipL::mSc* sporulation reporters into the *pyrE* loci of WT, Δ*tcdR*, Δ*spo0A* (65), and Δ*rstA* and visualized their expression on sporulation media (SMC agar). As expected, the P*sipL::mNG* and P*sipL::mSc* fluorescence was only observed in the mother cell for all strains analyzed, and no signal above background was observed for Δ*spo0A* (49, 60) (Figs 5 **& S8**). While the signal for *PsipL::mSc* was much brighter than for P*sipL::mNG* (Figs 5 **& S8A**), the P*sipL::mNG* reporter was useful for detecting *sipL* gene expression in live, sporulating cells that had completed asymmetric division but not engulfment (**Fig S8A, yellow arrow**). Cells at this stage of sporulation stain brightly in the forespore with Hoechst because the nucleoid is concentrated in a small region (57). This morphological information is largely lost for the P*sipL::mSc* reporter because the fixation procedure used to enhance the mScarlet signal causes the chromosome to fragment and obscures the forespore nucleoid signal (**Fig S5**). In contrast, the P*sipL::mNG* reporter can be used with the membrane stain FM4-64 and Hoechst nucleoid stain (**Fig S8B**) to glean useful cytological information on sporulation stage (57) as well as additional cellular phenotypes (66). Unfortunately, the mScarlet reporter is not compatible with the FM4-64 stain, limiting the utility of this reporter in cytological profiling studies.

**Figure 5.**
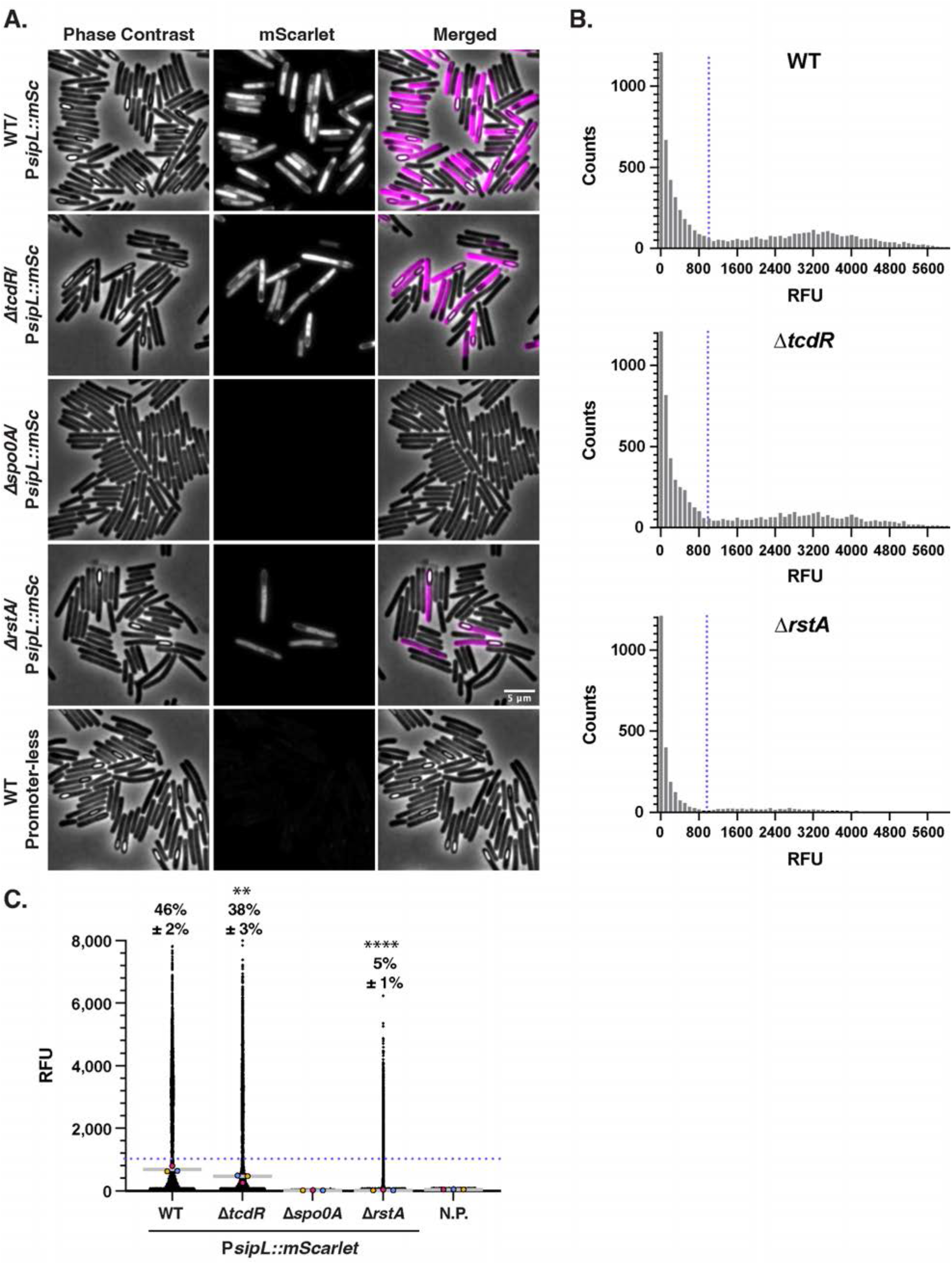
Sporulation gene expression is reduced in a toxin-less strain. (A) Fluorescence microscopy analyses of sporulating cultures of the indicated strains grown for 15 hrs on SMC sporulation agar followed by fixation. Phase contrast microscopy was used to visualize all bacterial cells. The merge of Hoechst (blue) and mScarlet (magenta) is shown. (B) SuperSegger-based quantification of the mean fluorescent intensity of each cell is shown as a black dot on the scatterplot. Individual cell intensities were quantified from three biological replicates, with at least two fields of view per strain per replicate. The magenta, yellow, and blue dots represent the median intensity for the first, second, and third biological replicates, respectively. The gray line represents the mean value of each replicate’s median value. N.R. indicates a strain harboring *mScarlet* with no upstream promoter region integrated into the *pyrE* locus. Percentage “Sporulation-ON” is displayed ± the standard deviation and was calculated using 1000 RFU signal as a cutoff (value displayed as a blue dotted line). This cut-off was determined using histogram analyses (Fig 5). A minority of points (< 1%) are outside the limits of the scatterplot; axes were adjusted to provide an optimal view of the scatterplot trends. Statistical significance was determined relative to wild type using a one-way ANOVA and Tukey’s test. *p<0.1; ** p < 0.01. (C) Histogram analysis of mean single-cell fluorescent intensities for the P*sipL::mSc* reporter in wildtype, *ΔtcdR* and Δ*rstA* demonstrates a bimodal distribution of cells as “Sporulation-ON” vs. “Sporulation-OFF”. Blue dotted line indicates the determined cutoff value of 1000 RFU which was also confirmed by visual inspection of phase-contrast images. Plotted data is a compilation of three biological replicates.

Despite these advantages, the P*sipL::mNG* reporter was too dim to be reliably quantified above *C. difficile*’s autofluorescence (data not shown). Quantification of the P*sipL::mSc* reporter allowed us to visualize two distinct populations of sporulating cells and non-sporulating cells (Fig 5A & 5B). A similar distribution of sporulating cells (population to the right of the dashed line) was observed for the WT and Δ*tcdR* P*sipL::mSc* reporter strains, whereas far fewer cells were found to be sporulating in the Δ*rstA* reporter strain consistent with the reduced sporulation phenotype of a Δ*rstA* strain (Fig 5B). When we attempted to quantify the proportion of cells that were “Sporulation-ON” using the same method as the toxin reporter calculations, i.e. using the 99^th^ percentile of the *Δspo0A* signal as the cutoff, 100% of wild-type cells were determined to be sporulating. Since this was clearly not the case based on phase-contrast microscopy and the reporter signal, we defined the “Sporulation-ON” population based on histogram analyses (Fig 5B) and manual inspection of the fluorescent signal in visibly sporulating cells vs. non-sporulating cells. With this cut-off, 46% of wild-type and 38% of Δ*tcdR* cells were identified as “Sporulation-ON” (Fig 5C). This slight decrease in proportion was statistically significant (p < 0.005); a slight reduction in the median fluorescence of the population was also observed between Δ*tcdR* and WT. Notably, the magnitude of the P*sipL::mSc* signal at the single-cell level decreased in Δ*rstA* relative to WT, and only 5% of cells were found to be “Sporulation-ON” (9-fold decrease, p < 0.0001).

Our reporter data was largely consistent with our functional analyses of sporulation using an ethanol resistance assay (**Fig S4C**). Δ*tcdR* produced functional spores at wild-type levels, while Δ*rstA* produced ∼3-fold less spores. In general, we detected smaller differences in sporulation efficiency relative to prior studies, where a *tcdR* Targetron insertion mutant made ∼2-fold more spores than wild-type 630Δ*erm* (35) and an *rstA* Targetron insertion mutant made 7- to 23-fold fewer spores than wild-type 630Δ*erm* (34, 35, 67). However, since the ethanol resistance assay measures both sporulation and germination efficiency, this assay tends to be more variable than the reporter assays (**Fig S9**) in our hands. This difference in variability is reflected in the statistical significance of the results obtained for each assay, with the reduction in sporulation for Δ*tcdR* and *ΔrstA* mutants relative to wild type achieving statistical significance for the reporter analyses (Fig 5C) but not the ethanol resistance assays. Regardless, these analyses validate the utility of the P*sipL::mSc* reporter for detecting cells that have completed asymmetric division.

### Transmission and virulence gene expression exhibit a division of labor in certain growth conditions

Having validated the single reporters P*tcdA::mNG* and P*sipL::mSc*, we next constructed dual reporter strains expressing both P*tcdA::mNG* and P*sipL::mSc* in the WT, Δ*tcdR*, Δ*spo0A*, and Δ*tcdR* strain backgrounds. These dual reporter strains allow us to directly visualize at the single-cell level the extent to which sporulating subpopulations and toxin-expressing subpopulations overlap (or bifurcate) in a coordinated manner, or if the *tcdA* and *sipL* genes are heterogeneously expressed independently of the other. The reporters also allow us to simultaneously assess the effect of media and growth conditions on toxin and sporulation gene expression. To construct the chromosomal dual reporter strains, the P*sipL::mSc* reporter was first inserted downstream of the *sipL* locus using allelic exchange and then the P*tcdA::mNG* reporter was integrated into the *pyrE* locus. The magnitude and proportion of cells expressing toxin and sporulation genes at the single-cell level was quantified using the same cut-off values as defined for the single reporters, since the toxin and sporulation gene reporters were expressed at similar levels and proportions for the WT dual reporter strain as the single reporter strain.

When the P*tcdA::mNG*-P*sipL::mSc* dual reporter strains were grown overnight in TY broth, which is traditionally used to induce maximal toxin gene expression (19), 27 ± 5% of the wild-type population was solely “toxin-ON” and 21 ± 6% of the population was solely “Sporulation-ON” (Fig 6). Simultaneous expression of both *tcdA* and *sipL* was observed in 11 ± 3% of the population (Fig 6, blue arrows) such that 41 ± 6% of the overall bacterial population was “Toxin-ON” (similar to what we observed in the single reporter strains in TY broth, Fig 4). About one-third of the population was “Sporulation-ON,” indicating that a sizable proportion of cells induce sporulation genes in TY broth, a property that was not possible to accurately quantify in our earlier analyses with the single P*tcdA::mNG* toxin reporter (Fig 4). In TY broth, Δ*tcdR* induced the *sipL* reporter at low levels, with only 2 ± 1% of cells being identified as “Sporulation-ON” (Fig 6C). However, when we used the bimodal distribution of the P*sipL::mSc* reporter to set the cut-off for “Sporulation-ON” cells (500 RFUs), approximately ∼4% of Δ*tcdR* cells were identified as “Sporulation-ON” in TY broth cultures. Importantly, this proportion more accurately reflects the visual inspection of the fluorescence microscopy images (Fig 6).

**Figure 6.**
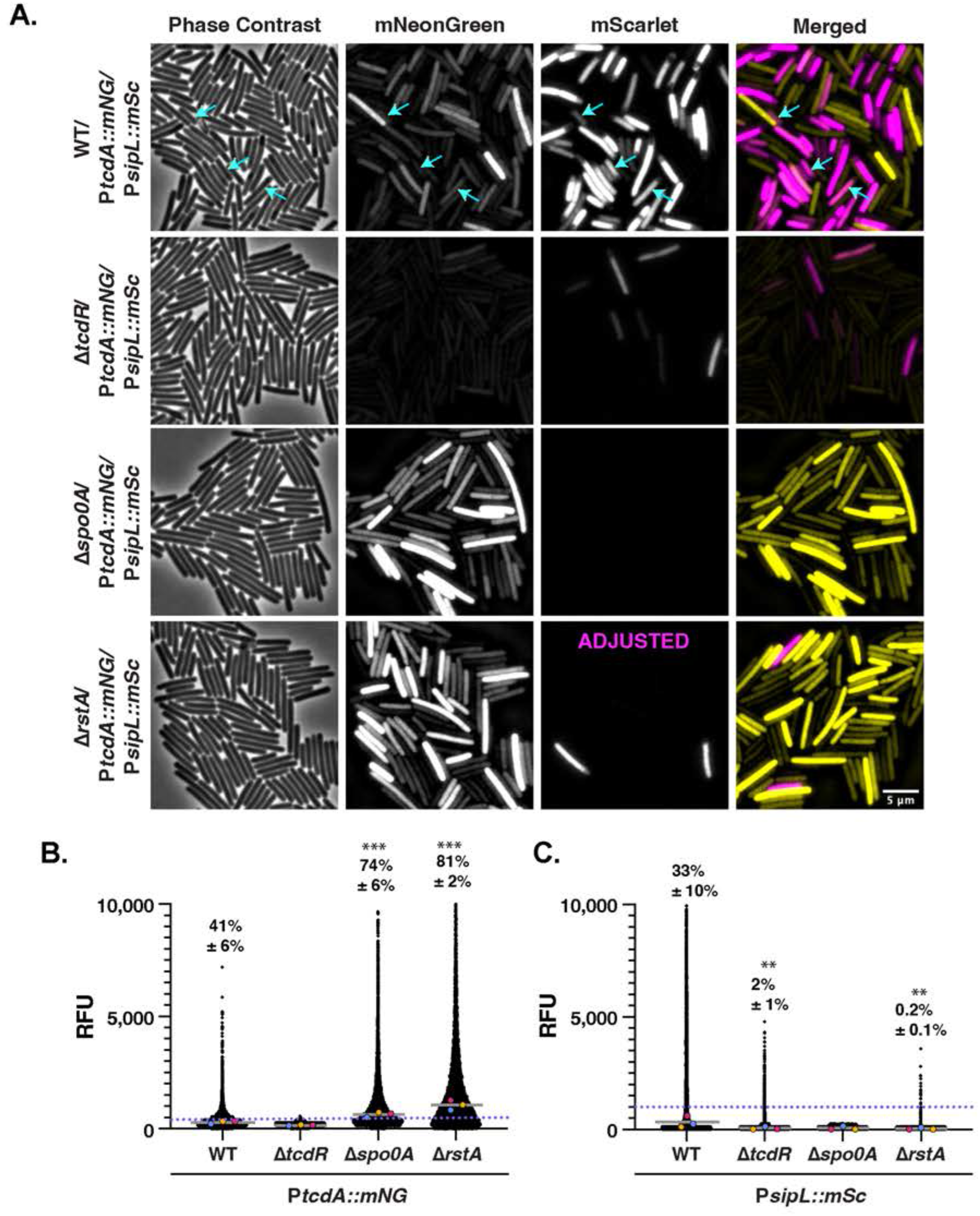
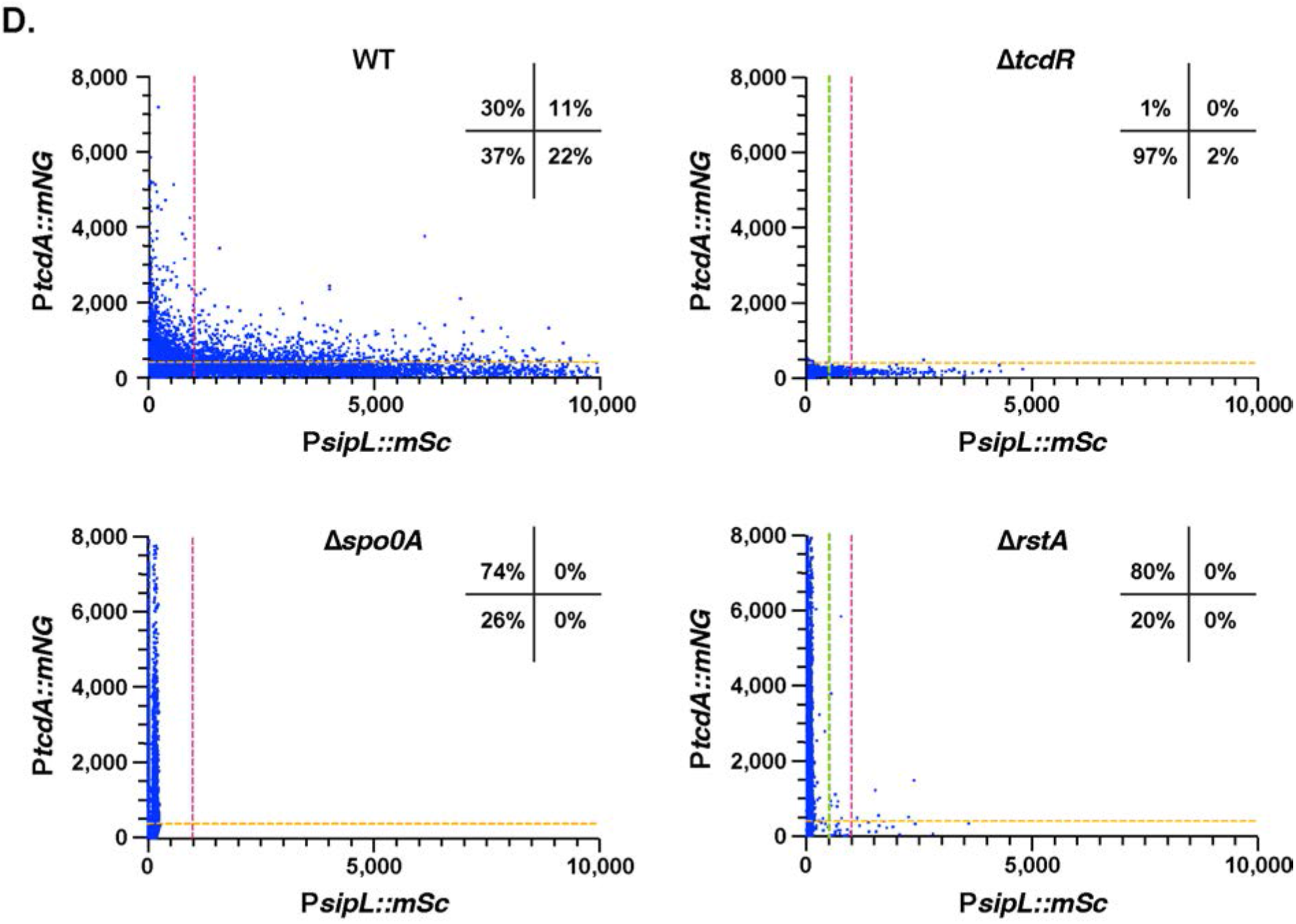
Simultaneous toxin and sporulation gene expression is observed in a subset of WT but not Δ*rstA* mutant cells during stationary phase growth in TY broth. (A) Fluorescence microscopy analyses of fixed bacterial cells after overnight growth in TY liquid media. Dual reporter strains contain P*sipL::mScarlet* and P*tcdA::mNeonGreen.* Dual reporter analyses were visualized in WT, Δ*tcdR*, Δ*spo0A* and Δ*rstA* strain backgrounds. Phase-contrast microscopy was used to visualize all bacterial cells. The merge of mNeonGreen (yellow) with mScarlet (magenta) signals are shown. Cells expressing both P*sipL::mScarlet* and P*tcdA::mNeonGreen* are highlighted with blue arrows. (B-C) SuperSegger-based quantification of P*tcdA::mNG* and P*sipL::mSc* reporters shows the mean fluorescent intensity of each cell as a black dot on the scatterplot for the indicated reporters. Individual cell intensities were quantified from three biological replicates, with at least two fields of view per strain per replicate. The magenta, yellow, and blue dots represent the median intensity for the first, second, and third biological replicates, respectively. The gray line represents the mean value of each replicate’s median value. Percentage “Toxin-ON” is displayed ± the standard deviation. “Toxin-ON” cells were calculated using the 99^th^ percentile of the Δ*tcdR* signal as a cutoff (value displayed as a pink dotted line). Percentage “Sporulation-ON” was calculated using 1000 RFU signal as a cutoff (value displayed as a pink dotted line; green dotted line represents the 500 RFU cut-off for “Sporulation-ON” cells for Δ*tcdR* and Δ*spo0A*). This cut-off was determined using the histogram analyses in Fig 5B. A minority of points (<1%) are outside the limits of the scatterplot; axes were adjusted to provide an optimal view of the scatterplot trends. Statistical significance was determined relative to wild type using a one-way ANOVA and Tukey’s test. ***p<0.001, ** p < 0.01. (D) Scatterplot analyses show single cell mean fluorescent intensities with P*sipL::mSc* on the x-axis and *PtcdA::mNG* on the y-axis. “Toxin-ON” cutoff is represented by the yellow dotted line which indicates 99^th^ percentile of the Δ*tcdR* signal. “Sporulation-ON” cutoff is represented by the magenta dotted line at 1000 RFU based on the histogram bimodal distribution analyses. Percentages indicate Toxin-ON/Sporulation-OFF in the top left quadrant, Toxin-ON/Sporulation-ON in the top right quadrant, Toxin-OFF/Sporulation-OFF in the bottom left quadrant and Toxin-OFF/Sporulation-ON in the bottom right quadrant. A minority of points (<1%) are outside the limits of the scatterplot; axes were adjusted to provide an optimal view of the scatterplot trends.

Consistent with our analyses of the single reporter P*tcdA::mSc* reporter in TY broth, the Δ*spo0A* and Δ*rstA* dual reporter strains induced the toxin reporter in a high proportion of cells (74 ± 6% and 81 ± 2%, respectively) and to higher levels than WT (Figs 4 & 6). Interestingly, only a small fraction of Δ*rstA* cells were identified as “Sporulation-ON” (0.2%). Taken together, these analyses indicate that overnight TY broth culture strongly induces both toxin and sporulation gene expression in wild-type cells. This result suggests that these two processes likely occur independent of the other, since similar proportions of cells appear to express both toxin and sporulation genes. Conversely, in Δ*rstA* a more prominent “division of labor” between these processes was observed, with cells primarily expressing *tcdA* often to high levels and a small subset inducing *sipL*. Notably, there was little overlap between these two sub-populations in contrast with WT.

Since toxin and sporulation gene expression are highly responsive to environmental conditions, we tested how different growth conditions affected their expression at the single-cell level. As a comparison to the TY liquid media tested above, we analyzed the behavior of the dual reporter strains on TY plates (Fig 7). Even though TY broth is used to induce toxin gene expression (19), when wild-type cells were growth on TY agar, they were 5-fold less likely to induce toxin (9% vs. 41% of the population was “Toxin-ON” on TY agar vs. TY broth), and the magnitude of toxin gene expression was also ∼2-fold lower on TY agar vs. TY broth (Figs 6 & 7). In contrast, wild-type cells were twice as likely to induce sporulation on TY plates (62% vs. 33% of cells were “Sporulation-ON” on TY agar vs. TY broth). Notably, when toxin gene expression was plotted against sporulation gene expression, minimal overlap was observed between the two populations for WT, with only 2 ± 0.1% simultaneously expressing toxin and sporulation genes when grown on TY plates compared to 11 ± 3% in TY broth.

**Figure 7.**
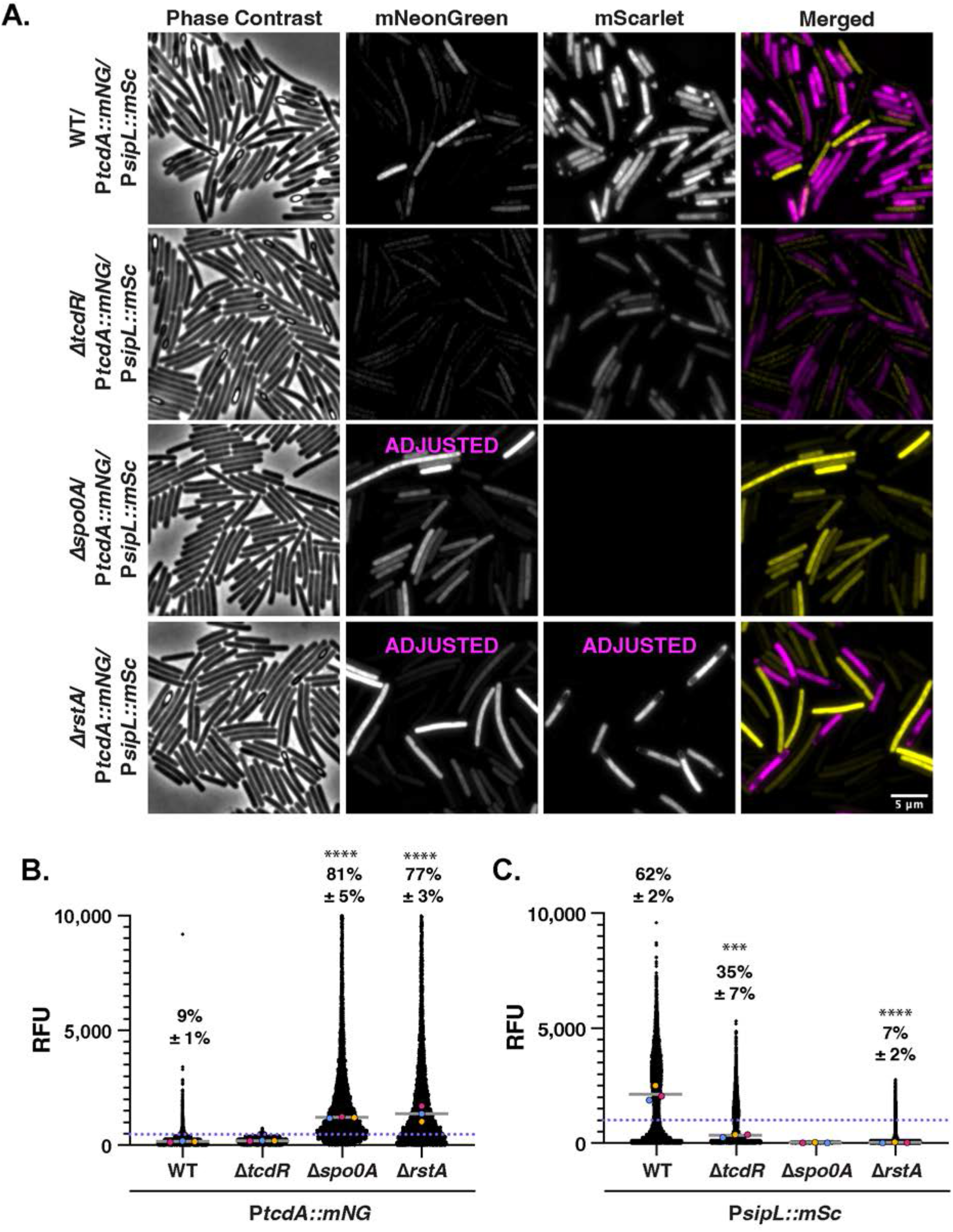
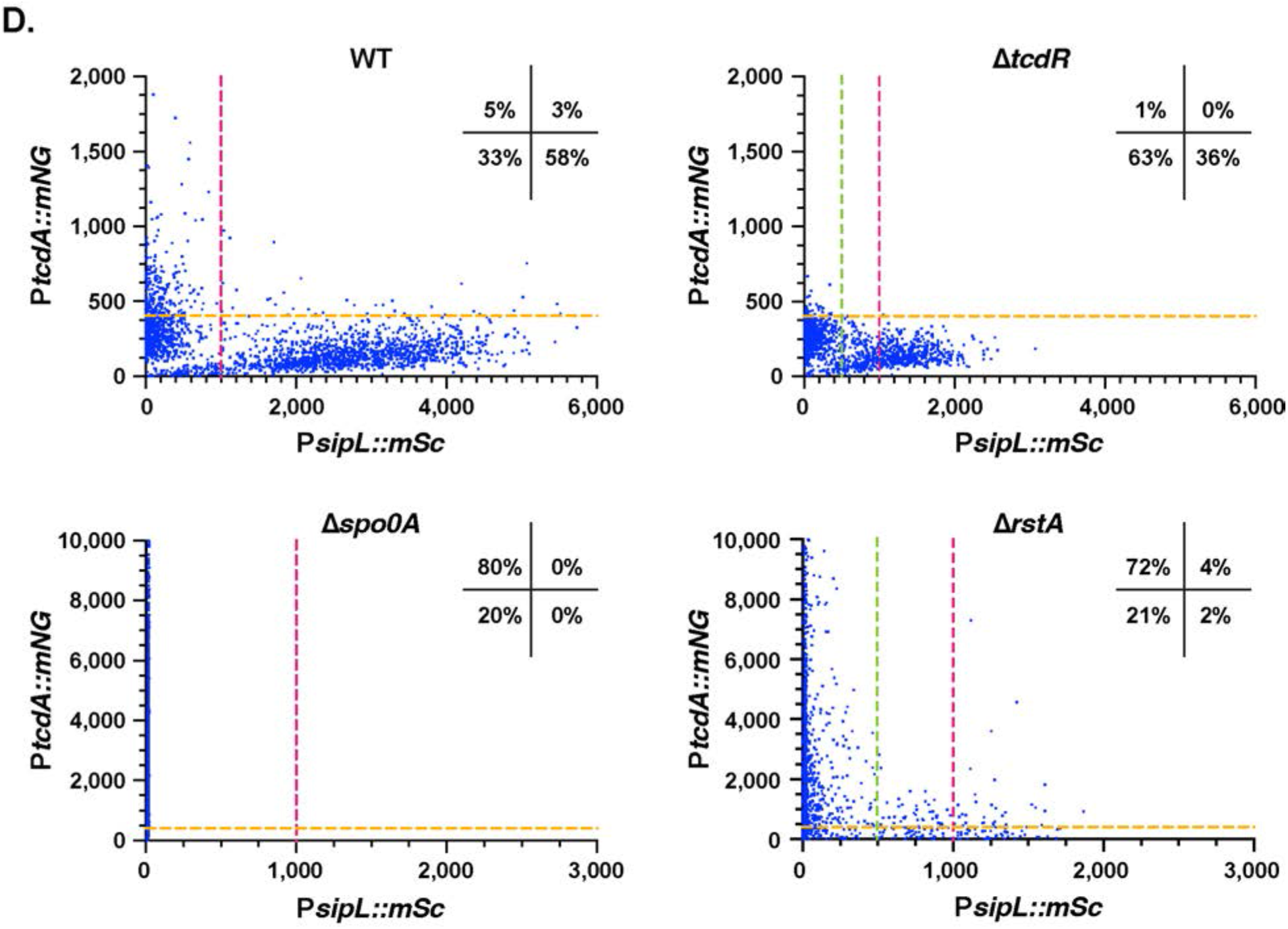
Minimal overlap between toxin and sporulation gene expression is observed during growth on TY plates. (A) Fluorescence microscopy analyses of fixed bacterial cells after growth on TY plates. Dual reporter strains contain P*sipL::mScarlet* and P*tcdA::mNeonGreen.* Dual reporter analyses were visualized in WT, Δ*tcdR*, Δ*spo0A* and Δ*rstA* strain backgrounds. Phase contrast microscopy was used to visualize all bacterial cells. The merge of mNeonGreen (yellow) with mScarlet (magenta) signals are shown. (B-C) SuperSegger-based quantification of P*tcdA::mNG* and P*sipL::mSc* reporters shows the mean fluorescent intensity of each cell as a black dot on the scatterplot for the indicated reporters. Individual cell intensities were quantified from three biological replicates, with at least two fields of view per strain per replicate. The magenta, yellow, and blue dots represent the median intensity for the first, second, and third biological replicates, respectively. The gray line represents the mean value of each replicate’s median value. Percentage “Toxin-ON” is displayed ± the standard deviation. “Toxin-ON” cells were calculated using the 99^th^ percentile of the Δ*tcdR* signal as a cutoff (value displayed as a blue dotted line). Percentage “Sporulation-ON” was calculated using 1000 RFU signal as a cutoff (value displayed as a pink dotted line; green dotted line represents the 500 RFU cut-off for “Sporulation-ON” cells for Δ*tcdR* and Δ*spo0A*). This cut-off was determined using the histogram analyses in Fig 5B. A minority of points (<1%) are outside the limits of the scatterplot; axes were adjusted to provide an optimal view of the scatterplot trends. Statistical significance was determined relative to wild type using a one-way ANOVA and Tukey’s test. ***p<0.001, ** p < 0.01. (D) Scatterplot analyses show single cell mean fluorescent intensities with P*sipL::mSc* on the x-axis and *PtcdA::mNG* on the y-axis. “Toxin-ON” cutoff is represented by the yellow dotted line which indicates 99^th^ percentile of the Δ*tcdR* signal. “Sporulation-ON” cutoff is represented by the pink dotted line at 1000 RFU based on the histogram bimodal distribution analyses. Percentages indicate Toxin-ON/Sporulation-OFF in the top left quadrant, Toxin-ON/Sporulation-ON in the top right quadrant, Toxin-OFF/Sporulation-OFF in the bottom left quadrant and Toxin-OFF/Sporulation-ON in the bottom right quadrant. A minority of points (<1%) are outside the limits of the scatterplot; axes were adjusted to provide an optimal view of the scatterplot trends.

While growth on TY agar vs. TY broth promoted sporulation by ∼3-fold in Δ*tcdR* and Δ*rstA* dual reporter strains (35% vs. 11% for Δ*tcdR* and 6% vs. 3% for Δ*rstA*), sporulation in these mutants was still reduced relative to WT, with Δ*tcdR* and Δ*rstA* inducing sporulation at ∼2- and ∼10-fold lower levels than WT on TY agar, respectively. Interestingly, Δ*tcdR* and Δ*rstA* strains also appeared to bifurcate into two distinct sub-populations of toxin and sporulation gene-expressing cells when grown on TY agar vs. in TY broth in the scatter plot analyses, suggesting that growth on a solid surface may promote a division of labor of toxin vs. sporulating gene expression for *C. difficile*. Notably, growth on TY agar did not change the frequency or magnitude of population-wide “Toxin-ON” cells detected for Δ*spo0A* and Δ*rstA* cells, with ∼80% of Δ*spo0A* and Δ*rstA* cells being designated “Toxin-ON” when grown on TY agar or plates. Similar trends in terms of the bifurcation in toxin and sporulation gene expression were observed for WT and Δ*tcdR* when the dual reporter strains were grown on SMC sporulation agar (**Figs S12-S13**), although this growth condition decreased toxin gene expression frequencies and levels for WT, Δ*spo0A*, and Δ*rstA* strains even further relative to TY agar.

Interestingly, cells that express toxin genes at high levels were less likely to express sporulation genes and vice versa such that cells that express both toxin and sporulation genes tend to express these genes at low to intermediate levels. This property is most evident in TY broth culture where wild-type cells were most likely to be simultaneously expressing toxin and sporulation genes. Taken together, our findings indicate that growth on agar plates promote a “division of labor” for *C. difficile*, where specific subsets of cells specialize into toxin vs. sporulation gene-expressing cells.

## DISCUSSION

Toxin production and spore formation are critical for *C. difficile* to cause and transmit disease, respectively, and several regulators that control both these important processes have been identified. While toxin production and spore formation are tightly coordinated in some organisms (10–15), the extent to which *C. difficile* coordinates these processes remained unclear in the absence of methods for simultaneously visualizing their induction at the single-cell level. In this paper, we address this question by developing a method for visualizing the expression of two genes of interest using chromosomally-encoded mNeonGreen and mScarlet transcriptional reporters in *C. difficile.* By coupling our dual reporter system with an automated method for quantifying the expression of these reporters at the single-cell level, we determined that toxin gene expression is generally lower in cells that are sporulating. Mutants that either cannot sporulate (Δ*spo0A*) or sporulate at greatly reduced levels (Δ*rstA*) express toxin genes at higher levels on a per-cell basis and in a greater proportion of cells regardless of the growth conditions tested (Figs 4-7).

However, sporulation and toxin gene expression are not always reciprocally regulated because a strain that cannot express toxin genes (Δ*tcdR*) expresses sporulation genes at lower levels but with similar frequencies at the single-cell level as wild-type *C. difficile* (Figs 4-7). Furthermore, growth of *C. difficile* on agar plates promoted a “division of labor” between transmission and virulence gene expression even if the media typically induces toxin gene expression in broth culture (i.e. TY media, Figs 6, 7, and **S12**). Interestingly, loss of the RstA regulator in TY broth caused *C. difficile* to primarily express toxin genes, whereas on TY plates, a small subset of Δ*rstA* cells simultaneously expressed both toxin and sporulation genes (4%). How RstA differentially regulates toxin and sporulation gene expression in broth culture vs. growth on a surface remains unclear, but in liquid culture, RstA may function to keep a clonal population uncommitted to either sporulation or toxin expression. Regardless, our data indicates that *C. difficile* bifurcates into specialized spore-formers and toxin-expressers in certain environmental conditions at least for the 630Δ*erm* strain, although we acknowledge that toxin (18) and sporulation gene expression may be differentially regulated at the single-cell level (35) in other *C. difficile* strains.

While the physiological importance of this “division of labor” remains unclear, it can be beneficial for individuals within a bacterial population to specialize and perform specific tasks that benefit the population as a whole (68). This phenomenon has been well-documented during biofilm formation (69), where metabolically intensive gene products are produced as “public goods”. For example, *B. subtilis* biofilms share energy-expensive matrix components to proximal cells (69), and distinct localizations for motile, matrix-producing cells versus sporulating cells are observed (70). Since *C. difficile* toxin is a metabolically costly secreted product, it may also be utilized as a “public good.” Indeed, with our toxin gene reporter, we observed that some *C. difficile* cells express toxin genes at extremely high levels relative to the rest of the population (Fig 4), and these cells appear unlikely to also express the sporulation reporter regardless of growth condition (Figs 6-7). Thus, *C. difficile* may use a similar strategy to differentiate into high toxin-producers so that other cells to focus their energy on sporulation or other tasks. Nevertheless, a critical question that arises from these analyses in laboratory media, is whether *C. difficile* employs this strategy during infection. During growth in the murine gut, a sub-population of *C. difficile* are sporulating, since ∼10% of *C. difficile* cells detected in the gut are present in the spore form (71). However, currently technologies are lacking to sufficiently analyze gene expression at the single cell level *in vivo*.

Notably, the chromosomal sporulation and toxin gene reporters we have developed could be useful for addressing these questions in the future, especially since our single-cell analyses of toxin and sporulation gene expression in different growth conditions highlights the complex intricacies of transmission and virulence gene expression. Future studies aimed at investigating how virulence and transmission gene expression are regulated in the host during infection will be crucial for characterizing *C. difficile* host-pathogen interactions. Does a “division of labor” between sporulation and toxin-expression occur in the host? Are toxin-expressing and sporulating subpopulations geographically distinct or stochastically overlapping? Where are transmission and virulence genes expressed and is there a pattern to their distribution? Utilizing these novel reporters in the context of an infection model could begin to address these important questions and gain mechanistic insight into *C. difficile* infection.

Finally, the chromosomally-encoded dual reporter system we have developed opens up numerous possibilities for deciphering the relationship between additional important processes in *C. difficile*. Recent work has revealed that *C. difficile* generates phenotypically heterogeneous sub-populations through multiple mechanisms, from bistability in toxin production (18); phase-variation in the expression of flagellar genes (40), genes encoding phage receptors (41), and two-component systems(72); as well as the expression of sigma factors that respond to stress (42). By optimizing the use of a transcriptional reporter that can overcome *C. difficile*’s natural autofluorescence (provided it is expressed at moderately high levels), it will now be possible to visualize how *C. difficile* specializes into different sub-populations and how nutritional inputs or growth conditions affect these different sub-populations. For example, our reporter system could be adapted to visualize how the levels of the important second messenger cyclic di-GMP affects flagella, toxin, biofilm production, and sporulation (20, 21, 40, 72, 73). These types of dual reporter analyses will undoubtedly provide important insight into the factors and mechanisms that control phenotypic heterogeneity in *C. difficile*.

## MATERIALS AND METHODS

### Bacterial strains and growth conditions

All *C. difficile* strains used for this study are listed in in Table S1 in the supplementary materials. All constructed strains derive from the sequenced clinical isolate 630, but the erythromycin-sensitive 630Δ*ermΔpyrE* is the parental strain used for all strain construction using *pyrE*-based allele-coupled exchange (ACE) (54). Strains were grown on brain heart infusion supplemented (BHIS) with yeast extract and cysteine (74), taurocholate (TA; 0.1% [wt/vol]; 1.9 mM), cefoxitin (8 μg/ml), thiamphenicol (10-15 µg/mL), and kanamycin (50 μg/ml) as needed. For ACE, the *C. difficile*-defined medium (CDDM) (75) was supplemented with 5-fluoroorotic acid (5-FOA; at 2 mg/ml) and uracil (at 5 μg/ml).

All *C. difficile* strains used for microscopy assays were first grown overnight from glycerol stocks on BHIS plates supplemented with TA (0.1% [wt/vol]). For broth culture microscopy assays, overnight cultures were inoculated from the BHIS-TA plates and grown in either BHIS or tryptone yeast broth (TY) (76). For plate-based microscopy assays, *C. difficile* strains were inoculated from BHIS-TA plates into BHIS liquid medium and grown until they were turbid. The cultures were then back-diluted 1:25 until they reached an optical density at 600 nm (OD_600_) between 0.4 and 0.7. 120 μL of this mid-log culture was used to inoculate either TY plates or SMC plates (77) for 18 hours prior to imaging.

*Escherichia coli* strains used for HB101/pRK24-based conjugations are listed in Table S1. *E. coli* strains were grown at 37°C with shaking at 225 rpm in Luria-Bertani (LB) broth. The medium was supplemented with ampicillin (50 μg/ml) and chloramphenicol (20 μg/ml) as needed.

### *E. coli* strain construction

All primers used for cloning are listed in Table S2 and plasmid maps for each construct are provided in Table S2. All plasmid constructs were sequence-confirmed using sanger sequencing through Genewiz. Plasmids were transformed into HB101/pRK24 *E. coli* and subsequently used to conjugate sequence-confirmed plasmids into *C. difficile*.

### *C. difficile* strain construction

Δ*tcdR* and Δ*rstA* deletion strains were generated using allele-coupled exchange (ACE) (54) with pMTL-YN3-Δ*tcdR* and pMTL-YN3-Δ*rstA*, respectively, and the parental 630Δ*erm*Δ*pyrE* strain. Single reporter strains and complementation strains were generated as previously described by conjugating HB101/pRK24-carrying pMTL-YN1C plasmids into Δ*pyrE*-based strains (65) using ACE. To generate dual reporter strains, P*sipL::mScarlet* was introduced downstream of the *sipL* locus of 630Δ*erm*Δ*pyrE,* 630Δ*ermΔtcdR*Δ*pyrE,* 630Δ*ermΔspo0A*Δ*pyrE* and 630Δ*ermΔrstA*Δ*pyrE* using ACE and pMTL-YN3-P*sipL::mScarlet*. The second reporter, P*tcdA::mNeonGreen*, was then introduced into the *pyrE* locus of the resulting strains using pMTL-YN1C-P*tcdA::mNeonGreen*. At least two clones of each strain generated by allelic exchange were phenotypically characterized prior to restoring the *pyrE* locus using pMTL-YN1C. At least two clones of every complementation strain were integrated into the *pyrE* locus and phenotypically characterized.

### Growth curves

*C. difficile* cultures were grown in 2 mL BHIS to stationary phase (4-5hrs) and then were back-diluted 1:25 in 2.5 mL BHIS until an OD_600_ of 0.5 was obtained. All strains were normalized to OD_600_ 0.5 if growth rates varied. Approximately 3.5 x 10^5^ CFU were inoculated into 150uL of the indicated media (∼50 uL of the OD_600_ 0.5 culture into 150 uL). 150uL of each strain was added to a 96-well plate (in technical triplicate) alongside appropriate blanks, and the plates were sealed with clear, gas-permeable ELISA plate sealers (R&D Systems). Plates were read in Epoch Plate Reader (BioTek) in the anaerobic chamber with OD_600_ readings performed every 15 minutes after a 2-minute linear shake.

### mScarlet maturation assay

For bulk fluorophore maturation quantification, replicate 96-well plates were set up as described above for the growth curves. Sealed plates were incubated overnight gently shaking in the anaerobic chamber. After overnight growth, plates were removed from the chamber, the seal was removed to expose cells to oxygen and then the plates were read in a Synergy H1 (BioTek) plate reader under ambient conditions at 37°C, with readings performed every 4 minutes for 24 hours, with orbital shaking every 5 seconds. The mScarlet fluorophore was excited at 560 nm and its emission was detected at 600 nm.

### Fixation protocol

Cells were fixed as previously described (43, 78). In brief, a 500 µL aliquot of cells grown in TY broth (or ∼1/2 loop of cells grown on plates was resuspended in 500 µL TY broth) was added directly to a tube containing 120 µL of a 5X fixation cocktail (100 μL of 16% (wt/vol) paraformaldehyde aqueous solution (methanol-free, Alfa Aesar) and 20 μL of 1 M NaPO_4_ buffer (pH 7.4). The samples were mixed and incubated aerobically for 30 minutes at room temperature in the dark followed by 30 minutes on ice in the dark. The fixed cells were washed 3 times in PBS and resuspended in 500 µL-1mL of PBS (depending on density of culture). Cells were immobilized on agarose pads and slides were incubated at 37°C for at least 2 hours to allow for mScarlet fluorophore maturation.

### Fluorescence microscopy

Bacterial cells were immobilized by spotting 1 µL of bacterial culture onto a 1.5% agar pad. Slides were incubated at 37°C for adequate maturation time (> 2 hr for mScarlet for live cell imaging, Fig 2). Nucleoid was stained using Hoechst 33342 (15 μg/ml; Molecular Probes). All images were acquired using a Leica DMi8 Thunder Imager equipped with a HC PL APO 63x/1.4 NA phase-contrast oil immersion objective. Excitation light was generated by a Lumencor Spectra-X multi-LED light source with integrated excitation filters. For all fluorescent channels aside from YFP, an XLED-QP quadruple-band dichroic beam-splitter (Leica) was used (transmission: 415, 470, 570, 660nm) along with an external filter-wheel (Leica). Phase contrast images were taken with a 50 ms exposure time. Hoechst was excited at 395/40nm (15% intensity), 100ms exposure, and emitted light filtered using a 440/40nm emission filter (Leica). mScarlet was excited at 555/38nm (20% intensity), with 150ms exposure time, and emitted light filtered using a 590/50nm emission filter (Leica). mNeonGreen images were captured with a YFP filter set (Chroma) equipped with a 500/20nm excitation filter, 515nm dichroic, 535/30 emission filter, excitation light was generated using the 510nm LED-line (100% intensity) and 387ms exposure. Emitted and transmitted light was detected using a Leica DFC 9000 GTC sCMOS camera. 3–4-micron Z-stacks were taken for each strain with 0.213 µm z-slices. All strains for a given experiment (6-8 strains) were spotted and captured sequentially on the same agar pad and Leica Adaptive Focus Control hardware autofocus was used to maintain focus at each position throughout the acquisition.

### Image analysis and quantification

After image acquisition, images were processed using the Leica “Instant Computational Clearing” (ICC) applied to the fluorescent channels to avoid bleed through fluorescent signal into neighboring cells. The adaptive strategy was run with the feature scale set to 2683nm and 98% strength. Images following ICC, were exported from the LASX software (Leica) and further processed using FIJI. Following export, images were cropped to remove out of focus regions and the best focused Z-plane was selected for each channel to correct for chromatic aberration using FIJI. Images were quantitatively analyzed using the SuperSegger pipeline (55) in MATLAB with the supplied ‘60x’ analysis settings. The output clist matrices containing all single-cell data were exported from MATLAB and the mean intensity was plotted as a scatterplot and/or histogram using Graphpad Prism (Version 9.0.2). Images were also analyzed manually in FIJI by examining the fluorescent intensities of visibly sporulating cells to confirm the cut-offs identified based on histogram analyses for “Sporulation-ON” were consistent with this visual inspection. Image scaling was adjusted to improve brightness and contrast for display and was applied equally to all strains in each experiment unless otherwise denoted by “ADJUSTED” on the displayed image. While some images are displayed with some saturation in the signal (clipping), no clipping occurred during image acquisition or quantification. At least three images per strain were captured in each replicate and every strain was analyzed with three biological replicates. Image analysis was performed on at least two positions per replicate. Statistical significance was determined using one-way ANOVA and Tukey’s test comparing the mean value of each replicate’s median value of three biological replicates (79).

### Ethanol-resistance sporulation assay

*C. difficile* cultures were grown in BHIS for 3-4 hrs, back-diluted 1:25, and after an OD_600_ of 0.35 – 0.7 was obtained, 120 µL of the log-phase culture was plated on 70:30 sporulation agar to form a lawn (77). Ethanol resistance was used to determine the sporulation efficiency of a given strain based on previously described procedures (80, 81). Specifically, after 24 hrs on 70:30 media, cells were resuspended in 70:30 broth to an OD_600_ of 1.0. Cells were immediately serially diluted in 70:30 broth and plated on BHIS + 0.1% taurocholate plates to enumerate all viable vegetative cells and spores. A 0.5 ml aliquot of culture was removed from the chamber, mixed with 0.5 ml 95% ethanol, vortexed and incubated at room temperature for 15 min, serially diluted in 1X PBS and plated on BHIS + 0.1% taurocholate plates to enumerate spores. After 24 hrs growth, CFU were enumerated, and the sporulation frequency was calculated as the number of ethanol-resistant spores divided by the total number of viable cells. The average ratio of ethanol-resistant colony forming units obtained from functional spores for a given strain relative to the average ratio determined for the wild type based was determined from a minimum of three biological replicates. A *spo0A* mutant was used as a negative control. Statistical significance was determined using one-way ANOVA and Tukey’s test.

### Western blotting

Samples for Western blot analyses were prepared as previously described (77), with minor adjustments. *C. difficile* was grown overnight for 18hrs in TY broth to induce toxin expression. For each strain 1mL of culture was pelleted, resuspended in 50 µL of PBS and samples were freeze-thawed for three cycles prior to resuspension in 100 µL of EBB buffer (8 M urea, 2 M thiourea, 4% [wt/vol] SDS, 2% [vol/vol] β-mercaptoethanol). The samples were boiled for 20 mins, vortexed, pelleted at high speed and resuspended in the existing buffer to maximize protein solubilization. Finally, the samples were boiled again for 5 minutes and pelleted at high speed and FSB was added. Samples were resolved on 7.5% SDS-PAGE gels and then transferred to an Immonilon-FL polyvinylidene difluoride (PVDF) membrane. The membranes were blocked in Odyssey blocking buffer with 0.1% (vol/vol) Tween20. Mouse anti-tcdA (REF) antibody was used at a 1:1000 dilution and chicken anti-GDH antibody was used at a 1:5000 dilution as a loading control. IRDye 680CW and 800CW infrared dye-conjugated secondary antibodies were used at a 1:20,000 dilution, and blots were imaged on an Odyssey LiCor CLx imaging system.

### Quantification of Western blotting

TcdA levels from Western blot analyses of three biological replicates were quantified using LiCor ImageStudio software and normalized to WT using the sum of all data points method(82). Statistical significance was determined using one-way ANOVA and Tukey’s test.

